# Short-term but not long-term triiodothyronine treatment improved cardiac function after myocardial infarction in male wild-type mice

**DOI:** 10.64898/2025.12.17.694596

**Authors:** Susanne C. Grund, Stefanie Dörr, G. Sebastian Hönes, Adrian D. Prinz, Christina Wenzek, Johannes Köster, Lars C. Moeller, Kristina Lorenz, Dagmar Führer

## Abstract

**Objectives:** Thyroid hormone (TH), especially triiodothyronine (T3), plays an important role in cardiac physiology and in the remodeling process following myocardial infarction (MI). We investigated the effects of short-term (until day 5) and long-term (until day 56) post-MI T3 treatment on cardiac function, infarct size, hypertrophy, and gene expression in mice without and with deletion of TRα, the main cardiac thyroid hormone receptor (wild-type (WT) and TRαKO, respectively)

**Methods:** WT and TRαKO mice underwent permanent left anterior descending coronary artery (LAD) ligation or sham surgery followed by either short-term T3 for 5 days post-MI or long-term T3 until day 56, including a subgroup in which T3 treatment commenced 14 days post-MI; all groups were followed up for 4 weeks. T3 was delivered via drinking water at 500 ng/ml. Cardiac function was studied with echocardiography (ejection fraction, EF), infarct size by histology (Sirius red), heart weight normalized to tibia length, and transcriptomic profiling (RNA-seq) in WT hearts.

**Results:** Short-term T3 improved EF in WT but not in TRαKO mice without induction of hypertrophy or changes in infarct size in either genotype. Long-term T3 induced cardiac hypertrophy in both WT and TRαKO mice. However, long-term T3 did not improve EF or reduce infarct size. In TRαKO mice, baseline EF post-MI was preserved without T3, but T3 treatment decreased EF. RNA-seq in long-term treated WT mice suggested modulation of Rho-GTPase signaling, mitochondrial biogenesis, and immune activation by T3.

**Conclusions:** T3 therapy post-MI improved cardiac function only when applied acutely and for a short term. Long-term exposure led to cardiac hypertrophy without functional improvement and may even worsen cardiac function in TRα-deficient settings. Timing, duration, and receptor status are highly relevant for TH-based interventions in MI.

## Introduction

Cardiovascular diseases remain the leading cause of morbidity and mortality worldwide, with myocardial infarction (MI) being a major contributor. Beyond traditional risk factors such as hypertension, diabetes, and dyslipidemia, endocrine influences on cardiac health are increasingly recognized. Among these, thyroid hormones (TH) play a pivotal role in regulating cardiovascular physiology, yet their contribution to the risk, manifestation, and prognosis of MI remains incompletely understood.

Thyroid hormones exert wide-ranging effects on the cardiovascular system by modulating heart rate, contractility, vascular resistance, and myocardial oxygen demand. In states of hyperthyroidism, elevated triiodothyronine (T3) and thyroxine (T4) accelerate metabolism, increase sympathetic sensitivity, and enhance cardiac output. These changes predispose individuals to arrhythmias, left ventricular hypertrophy, and an increased risk of ischemic events due to heightened myocardial oxygen consumption (Kahaly & Dillmann, 2005; Klein & Ojamaa, 2001). Conversely, hypothyroidism is characterized by reduced cardiac output, bradycardia, impaired endothelial function, and elevated systemic vascular resistance.

Hypothyroidism is also linked to atherogenic lipid profiles, which may accelerate coronary artery disease and increase susceptibility to MI (Jabbar et al., 2017; Klein & Danzi, 2007). Particular attention has been given to low T3 syndrome, a transient reduction in circulating T3 levels observed in approximately 20–30% of patients hospitalized with acute MI (Iervasi et al., 2003; Li et al., 2011; Zhang et al., 2012). Several clinical studies have shown that reductions in T3, often accompanied by increased reverse T3 (rT3), correlate with worse short- and long-term outcomes, including impaired ventricular recovery and higher mortality (Friberg et al., 2001; Lymvaios et al., 2011; Zhang et al., 2012). However, findings across studies remain heterogeneous, partly due to variations in patient selection, timing of hormone measurement, and comorbid conditions (Eber et al., 1995; Friberg et al., 2002). Importantly, low T3 syndrome has been reproduced in animal models of MI and T3 supplementation improved cardiac function (Forini et al., 2014; Nicolini et al., 2016), suggesting both diagnostic and therapeutic relevance.

Thyroid hormone administration has emerged as a potential therapeutic approach after MI. Mechanistically, TH treatment reduced apoptosis, promoted angiogenesis, restored mitochondrial function, and activated cardioprotective signaling pathways (Chen et al., 2008; Forini et al., 2014; Mourouzis et al., 2013; Sabatino et al., 2016). Clinical studies remain limited and small in scale, but some reported improved ventricular performance and wall motion in MI patients treated with T3 (Pingitore et al., 2008; Pingitore et al., 2019). Ex vivo and in vivo data from rodent models showed that T3 treatment improved cardiac recovery and function after ischemia/reperfusion. Ex vivo, recovery of left ventricular developed pressure in isolated rat hearts subjected to ischemia/reperfusion was markedly improved when treated with T3 during reperfusion (Bi et al., 2019; Pantos et al., 2009). In vivo, T3 treatment post-MI seemed beneficial and successful in improving cardiac function when given as two single doses immediately and after 24 hours or for 3 to 12 days post-MI (Cerullo et al., 2025; Chen et al., 2008; de Castro et al., 2016). Given with a 72-hour delay but for 4 weeks from then on, T3 also improved cardiac function and EF (Forini et al., 2011).

These data suggest that T3 treatment improves post-MI outcome in patients. The clinically relevant question is when to start and which treatment duration improves post-MI cardiac function. Are an early start and short duration sufficient, as the ex vivo data indicated, or would a longer treatment duration induce physiological hypertrophy and improve cardiac function further?

Thyroid hormone receptor α (TRα) is the predominant isoform expressed in cardiomyocytes, whereas TRβ is more abundant in the liver and pituitary (Yen, 2001). While the effect of T3 treatment has been studied, the role of TRα in mediating cardiac outcomes in response to T3 treatment after MI and has not yet been specifically addressed. With the emergence of TR isoform-specific agonists in clinical practice (Harrison et al., 2024), another clinically relevant question is: is TRα the isoform to target post MI?

To address the role of TRα as well as duration and timing of T3 treatment in post-MI cardiac function, we investigated the effects of short- and long-term T3 treatment on cardiac function and hypertrophy after MI in wild-type (WT) and TRα knockout (TRαKO) mice.

## Material and Methods

### Animals and treatment

All animal experiments were approved by the Landesamt für Verbraucherschutz und Ernährung Nordrhein-Westfalen (LAVE-NRW, AZ: 81-02.04.2020.A282) and performed in accordance with the German regulations for Laboratory Animal Science and the European Health Law of the Federation of Laboratory Animal Science Associations. TRα knockout (TRα^0/0^) mice, referred to as TRαKO, were obtained from the European Mouse Mutant Archive (Gauthier et al., 1999; Gauthier et al., 2001).

Mice were housed in the central animal facility at the University Hospital Essen in individually ventilated cages at 21±1°C in an alternating 12:12-hour light-dark cycle and fed standard chow (Sniff, Soest, Germany) and tap water provided ad libitum.

Male wild-type (WT) and TRαKO mice (littermates) were studied. Mice were randomized to the following groups (n = 10/group): Sham-operated (Sham), Sham plus T3 treatment (Sham + T3), myocardial infarction by permanent ligation of the left anterior descending (LAD) coronary artery (MI), and MI plus T3 treatment (MI + T3). T3 was administered via drinking water at 500 ng/ml. Short-term treatment: 5 consecutive days post-MI. Long-term: 8 weeks of continuous treatment, or delayed start (day 14 post-MI).

### Myocardial Infarction Model

At the age of 8-12 weeks, the left anterior descending coronary artery of WT and TRαKO mice was permanently ligated to induce MI. After anesthesia with a combination of medetomidine (0.5 mg/kg), midazolam (5 mg/kg) and fentanyl (0.05 mg/kg) (i.p.), the chest was opened between the third and the fourth rib. LAD was occluded using an 8 − 0 silk suture (MI) or the suture was removed again (sham). Anesthesia was antagonized using a combination of atipamezole (0.75 mg/kg), flumazenil (0.5 mg/kg) and buprenorphine (0.1 mg/kg) (s.c.). For analgesia, animals received metamizole in drinking water (300 mg/kg, p.o.) starting two days before surgery up to 7 days after surgery.

### Echocardiography

Echocardiography was performed under pentobarbital anesthesia (20-35 mg/kg body weight, i.p.) using a Vevo^®^3100 high-resolution imaging system (FUJIFILM VisualSonics, Amsterdam, The Netherlands) and the MX550D transducer. Animals were kept on a heating plate to maintain body temperature throughout the procedure. Parasternal long axis images and the VisualSonics Cardiac Measurements software were used to evaluate the ejection fraction. The data represent the average of at least four cardiac cycles. The investigator was blinded regarding treatment groups during measurements and data analysis. Measurements at heart rates below 430 bpm were excluded (Brand et al., 2025). For short term T3 treatment experiments measurements were performed before and four weeks after LAD ligation or sham surgery. For the long term T3 treatment group echocardiography was performed before, 2 weeks, 4 weeks and 8 weeks after LAD or sham surgery.

### Histological staining

Hearts were fixed in 4% (m/V) paraformaldehyde and embedded in paraffin. Heart tissue was sliced transversely (5μm). Sections of the midpapillary region were stained with Sirius Red (Schmid et al., 2015; Tomasovic et al., 2020; Vettel et al., 2017). For the determination of the infarct size, the epicardial and endocardial infarct length and circumference was evaluated using ImageJ (Vettel et al., 2017). Infarct size was calculated using this equation: [(epicardial infarct length/epicardial circumference) + (endocardial infarct length/endocardial circumference)/2] * 100 13. Analysis and quantification were performed by blinded researchers.

### RNA Isolation and Gene Expression Analysis

Total RNA was isolated from homogenized heart tissue from the remote area. After preincubation of the homogenized heart with proteinase K for 20 min at 55 °C, RNA isolation was performed following the manufacturer’s instructions (RNeasy Mini Kit, QIAGEN).

Concentration and quality of RNA was measured with Agilent Bioanalyzer Nano (Agilent, Santa Clara, CA USA). RNA (900 ng) was reverse transcribed into cDNA with SuperScript III (Invitrogen) and random hexamers. Quantitative PCR (qPCR) was performed on Real-Time PCR-Cycler CFX384TM C100 Touch Thermal (Bio-Rad). Transcripts were quantified by Eva Green Mix (Bio-Rad). Relative mRNA levels were normalized to the expression of *Gapdh* mRNA and calculated with efficiency correction. The primers used to amplify THRB were 5’-GGA CAA GCA CCC ATC GTG AA-3’ (forward) and 5’-ACA TGG CAG CTC ACA AAA CAT-3’ (reverse).

For RNA-Sequencing, libraries were prepared with Lexogens QuantSeq 3’ mRNA-Seq Library Prep Kit FWD, quality controlled with Qubit (Invitrogen, Waltham, MA, USA) and library quant qPCR, and sequenced on a NextSeq500 75 cycles (Illumina, San Diego, CA, USA). Data were analyzed with the rna-seq-kallisto-sleuth workflow (https://github.com/snakemake-workflows/rna-seq-kallisto-sleuth). Specifically, sequences were trimmed with Cutadapt (Martin, 2011). Transcripts were quantified while accounting for 3’ mRNA-seq by mitigating the leaking of non-MANE transcript expression into the MANE transcript expression estimates. This was achieved by mapping reads to the whole transcriptome using BWA-MEM (Li, 2013) keeping only reads that mapped closest to the 3’ end of the MANE transcript of each gene. Then, Kallisto (Bray et al., 2016) was used to quantify the MANE transcripts with the filtered reads. Integrated normalization and transcript differential expression analysis was conducted with Sleuth (Pimentel et al., 2017), using the likelihood ratio test method while correcting for experimental cohort related batch effects.

Sleuth effect size estimates (beta-scores) were calibrated to represent log2-fold changes by transforming input counts x with log2(x + 0.5). False discovery rate was controlled with the Benjamini-Hochberg method (Pimentel et al., 2017). Genes were sorted in a threshold-free approach utilizing the pi-value (Xiao et al., 2014), which we reinterpreted to combine FDR with absolute values of sleuth beta-scores of the respective comparison variable, such that those genes with the strongest combination of significance and effect size are listed at the top. Gene ontology enrichment was performed on expressions of MANE transcripts (according to Ensembl Mus musculus release 111) using goatools (Klopfenstein et al., 2018), while defining transcripts with FDR<=0.05 as foreground. All results were visualized using Datavzrd (Wiegand et al., 2025) and Vega-Lite (Klopfenstein et al., 2018).

### Statistical Analysis

Results are mean ± SD. Two-way ANOVA (factors: genotype, treatment, MI status), with post-hoc tests. Significance p < 0.05.

## Results

### Acute T3 treatment after MI improves cardiac function in WT mice

In the short-term protocol (Fig. 1A), WT and TRαKO mice were treated with T3 or vehicle directly after MI induction by LAD ligation or sham surgery for 5 days. Cardiac effects of T3 were assessed at day 28. T3 treatment did not increase heart weight normalized to tibia length (HW/TL) in any of the short-term treated groups (both WT and TRαKO), relative to their respective controls (Fig. 1B). Without T3, however, the heart weight of vehicle treated TRαKO mice was reduced compared to vehicle treated WT mice (Fig. 1C). This difference was absent after MI.

**Figure 1.**
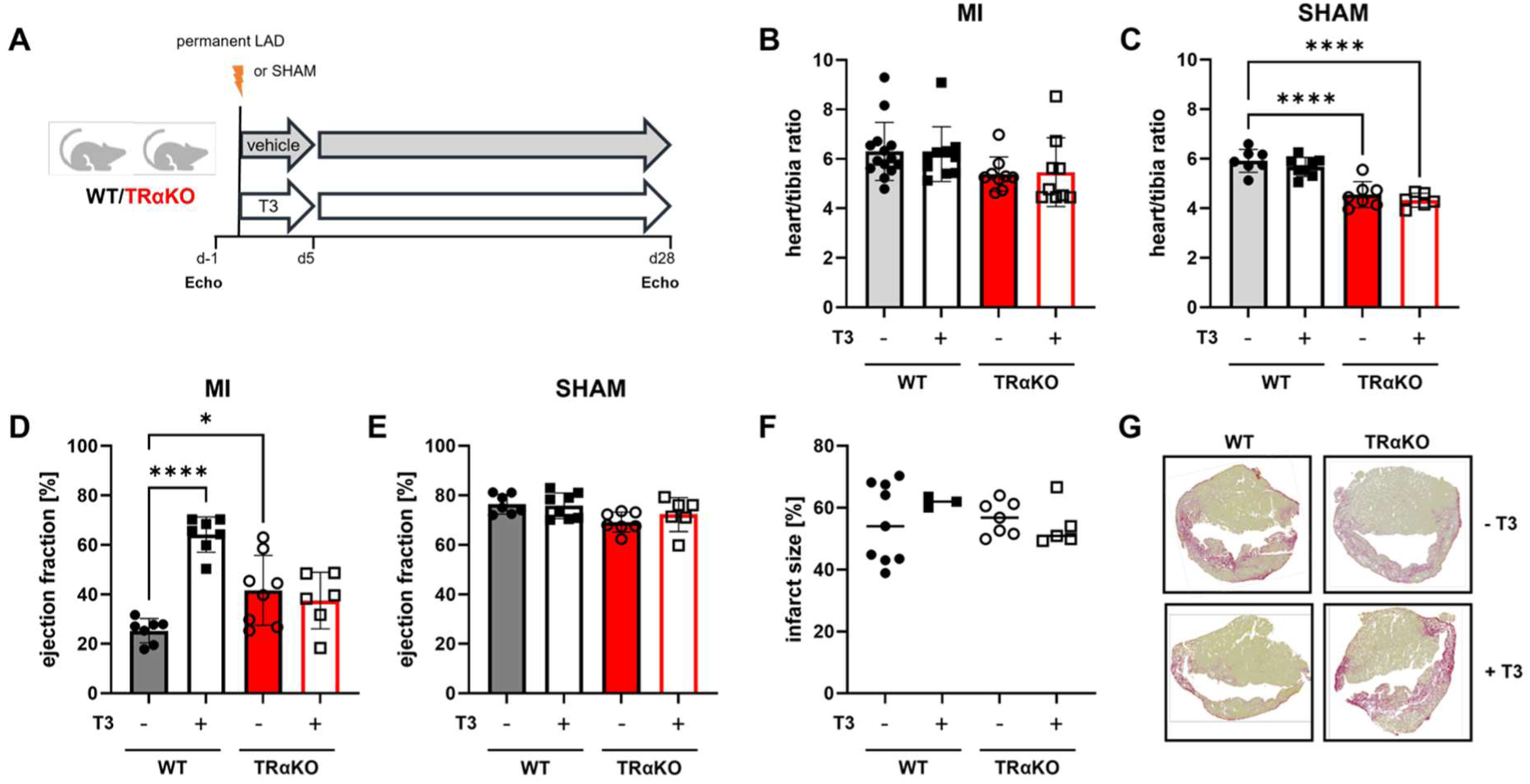
Acute T3 treatment improves functional outcome after MI in WT mice. (A) Overview: mice were treated with 500 ng/ml T3 or vehicle right after inducing permanent LAD for 5 days post MI. Echocardiography was performed before induction of MI (-1) and 14 and 28 days post MI. (B) Heart weight in WT and TRαKO mice with and without T3 treatment after dissection at day 28 post MI or (C) SHAM surgery (in relation to tibia length). (D) EF in WT and TRαKO MI and E) Sham mice with and without T3 treatment after dissection at day 28 post MI. (F) Infarct size EF in WT and TRαKO mice with and without T3 treatment after dissection at day 28 post MI. (G) Sirius red staining from exemplary WT and TRαKO hearts 28 days post MI. MW ± SD; *p≤0.05, ****p≤0.0001

Five days of T3 treatment following permanent LAD ligation led to an improved cardiac function of wild-type (WT) mice as assessed 4 weeks after MI (Fig. 1D): While the EF of vehicle treated WT mice was reduced to 25.32 ± 4.97 %, the EF in the WT MI + T3 group was preserved with 64.19 ± 7.13% (P<0.0001).

EF in vehicle treated TRαKO was less reduced compared to vehicle treated WT mice. In contrast to WT mice, TRαKO mice showed no difference in EF between the MI and MI + T3 groups in the short-term treatment arm (Fig. 1D), indicating that the benefit of T3 in improving systolic function depends on TRα. Assessment of infarct size using Sirius red staining revealed no significant differences between MI and MI + T3 groups in either genotype (Fig. 1E+F).

### Long-term T3 treatment leads to hypertrophy in MI and sham

In the long-term experiments (Fig. 2A), T3 treatment was started either directly after MI/sham surgery or post-MI at day 14 and continued until day 56.

**Figure 2.**
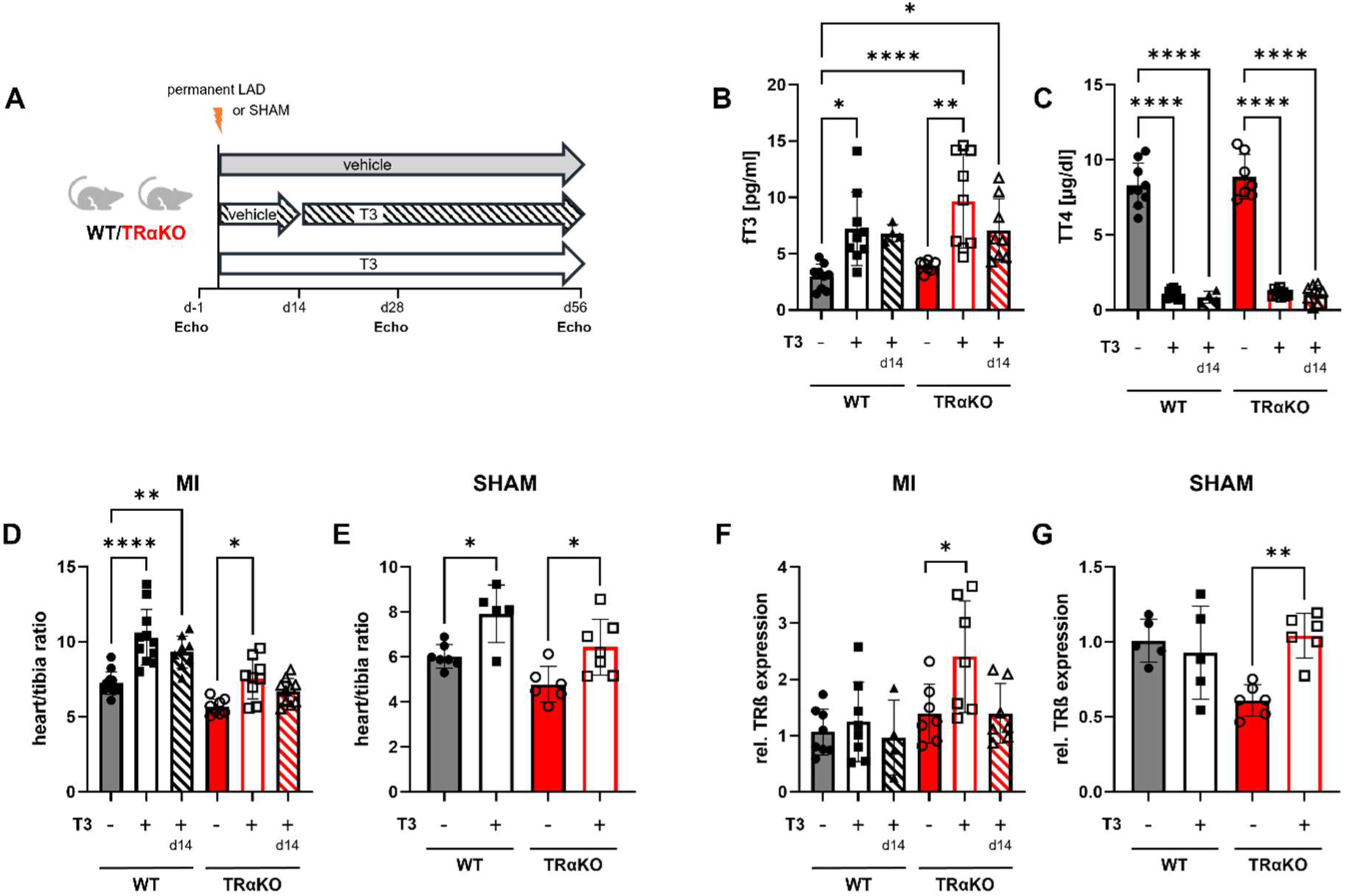
Long-term T3 administration increases heart weight. (A) mice were treated with 500 ng/ml T3 or vehicle directly after inducing permanent LAD or 14 days post LAD until day 56. Echocardiography was performed before induction of MI, 14-, 28- and 56-days post MI. (B) fT3 and TT4 serum concentrations. (C) Heart weight in relation to tibia length in WT and TRαKO mice after MI od SHAM. (F) TRβ mRNA expression in MI and (G) SHAM WT and TRαKO mice treated with T3 for 56 days. MW ± SD; *p≤0.05, **p≤0.01, ****p≤0.0001

T3 treatment after MI produced marked changes in systemic thyroid hormone levels both when initiated immediately after MI or later from day 14 onwards as assessed 8 weeks after surgery. In line with this, free T3 was elevated (Fig. 2B), while total T4 was suppressed (Fig. 2C) in WT and TRαKO mice.

Heart weight normalized to tibia length (HW/TL) was significantly increased in T3-treated WT mice, in both treatment groups, the ones treated with T3 for 8 weeks post-MI or for 6 weeks started 14 days post-MI (Fig. 2D). T3 treatment also increased heart weight in TRαKO mice, although to a lesser degree and only when treated for 8 weeks post-MI (Fig. 2D). A comparable T3 effect was seen after sham surgery of WT and TRαKO mice (Fig. 2E). Of note, we found an upregulation of TRβ mRNA expression in the TRαKO hearts with T3 treatment (Fig 2F+G).

### Long-term T3 treatment does not improve functional outcome after MI

In untreated WT mice, MI led to a severely impaired cardiac function with a reduction of EF to 37.92 ± 11.44% whereas Sham operated mice were unaffected (Fig. 3A). Neither an 8 week nor a 6 week T3 treatment (started directly or 2 weeks post-MI, respectively) led to significant improvement in EF compared to mice treated with vehicle.

**Figure 3.**
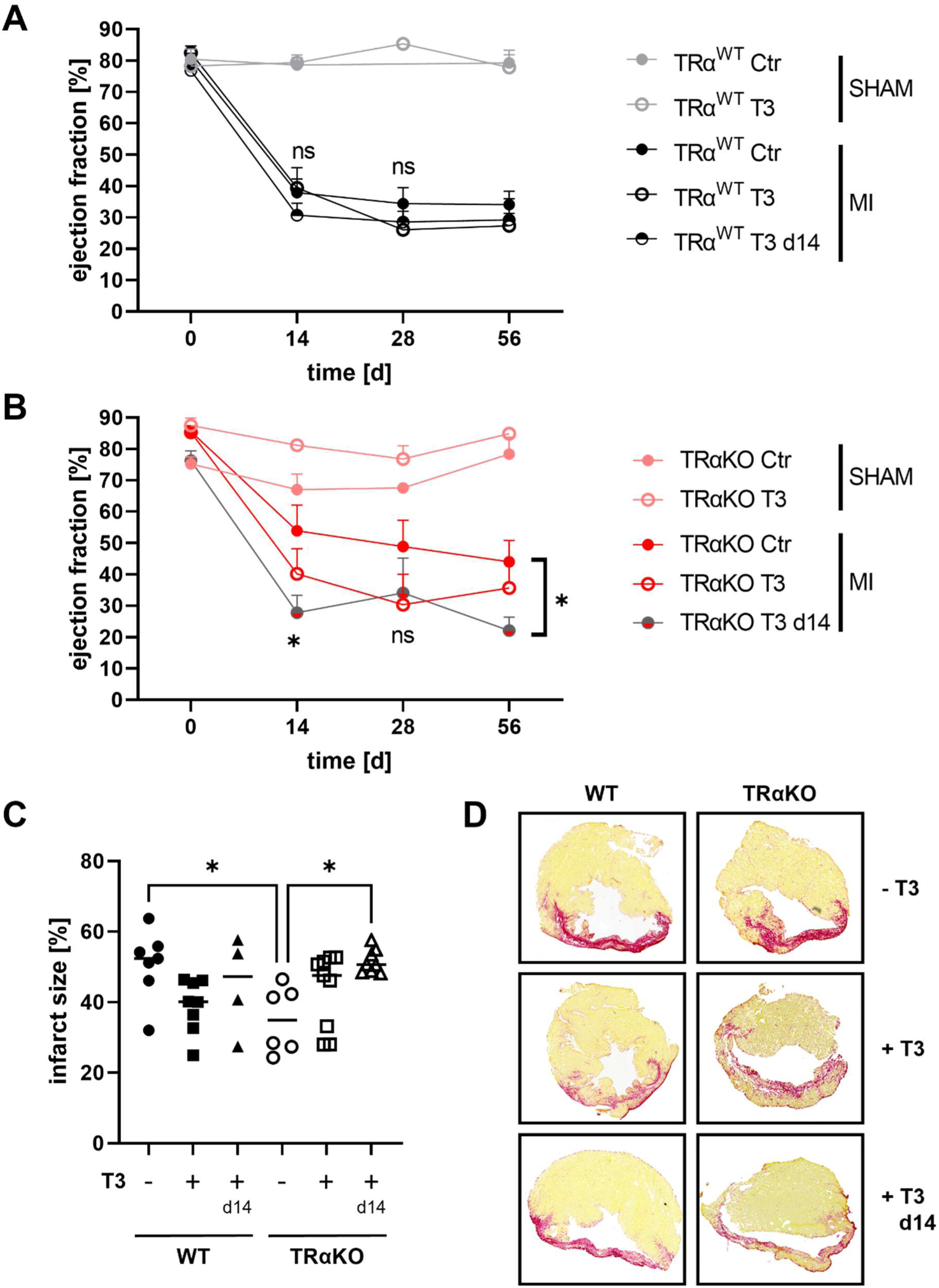
Long-term T3 treatment does not improve functional outcome after MI. (A) Cardiac function (EF) in WT mice after induction of MI or SHAM surgery, treated with T3 or vehicle for 56 days. (B) Cardiac function (EF) in TRαKO mice after induction of MI or SHAM surgery, treated with T3 or vehicle for 56 days. (C) Scar size determined by quantification of sirius red staining after T3 treatment or vehicle in WT and TRαKO mice (D) Exemplary WT and TRαKO hearts after T3 treatment either immediately or from day 14 post-MI or vehicle (Sirius red staining). MW ± SD; *p≤0.05

In vehicle treated TRαKO mice, baseline EF after MI was less severely affected compared to WT mice (Fig. 3B). However, as in WT mice, long-term T3 treatment did not result in an improved EF.

Infarct scar size 8 weeks post-MI was assessed by Sirius red staining, which revealed reduced scar size in vehicle treated TRαKO mice and – even though not significant – also in T3 treated WT mice (Fig. 3C,D).

### Transcriptomic analysis of regulated pathways in T3-treated vs. vehicle/control treated WT MI mice

RNA-seq and subsequent pathway analyses of differentially regulated genes of WT hearts following long-term T3 treatment after MI revealed several activated and inactivated regulatory mechanisms of heart function after MI (Table 1). Genes involved in Rho-GTPase signaling were significantly modulated by long-term T3 treatment in WT mice, namely their expression pattern suggested an overall inhibition of the Rho-GTPase pathway (Fig. 4A).

**Figure 4.**
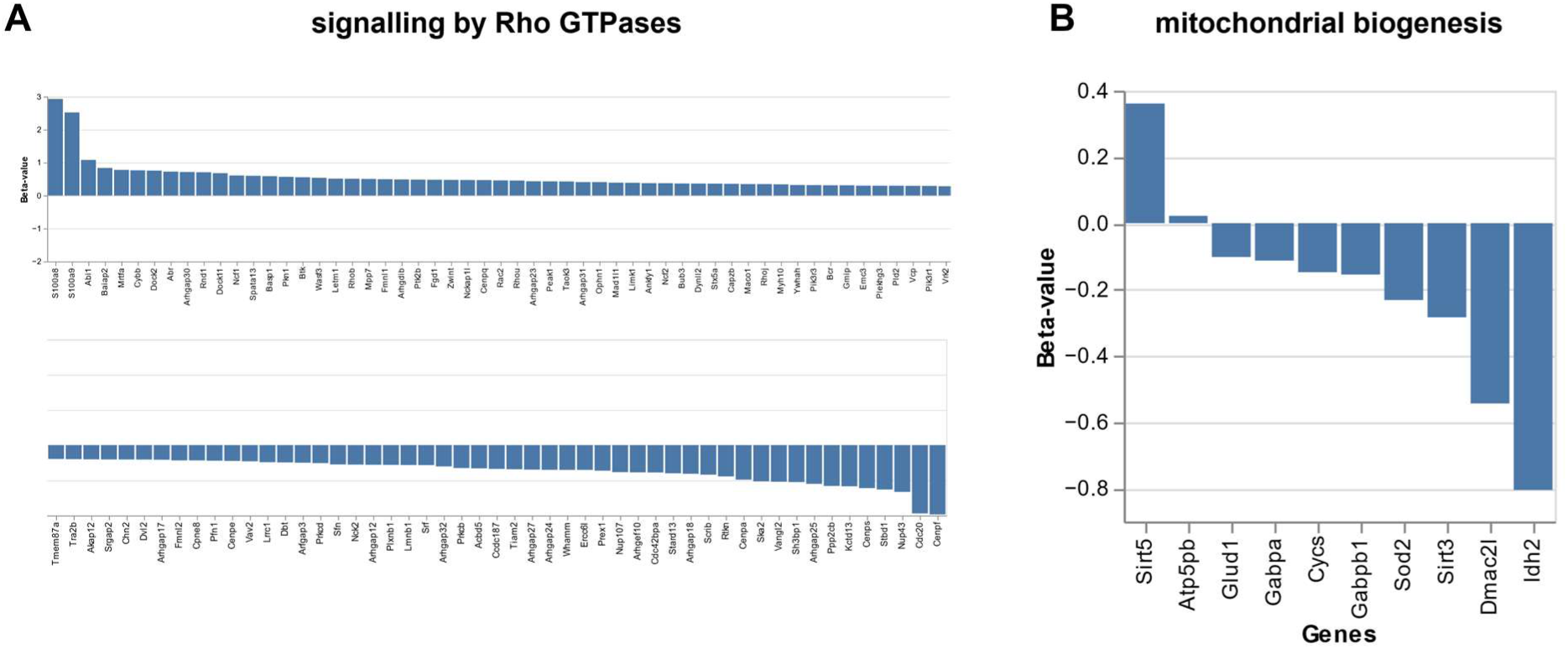
Differentially regulated genes associated with RHO-GTPase signaling pathway and mitochondrial biogenesis. Shown are waterfall plots of regulated genes of the ‘signaling by Rho GTPases’ (A) and ‘mitochondrial biogenesis’ pathways from Table 1.

**Table 1.** Pathway analysis of regulated genes in WT MI vs. WT MI+T3 hearts. Shown are the total perturbation accumulation, combined FDR values, inhibition or activation of pathway, gene ratio and the signed pi value.

| Name | total perturbation accumulation | Combined FDR | Status | gene ratio | signed pi value |
| --- | --- | --- | --- | --- | --- |
| Signaling by Rho GTPases, Miro GTPases and RHOBTB3 | -20.36 | 0.28 | Inhibited | 0.18% | -11.21 |
| Signaling by Rho GTPases | -20.36 | 0.28 | Inhibited | 0.18% | -11.21 |
| Cell Cycle, Mitotic | -9.63 | 0.28 | Inhibited | 0.20% | -5.30 |
| Cell Cycle | -9.13 | 0.28 | Inhibited | 0.17% | -5.03 |
| G alpha (q) signalling events | 3.27 | 0.09 | Activated | 1.13% | 3.39 |
| Mitotic Prometaphase | -2.63 | 0.24 | Inhibited | 0.57% | -1.65 |
| EPH-Ephrin signaling | -2.20 | 0.21 | Inhibited | 1.61% | -1.51 |
| RHO GTPase Effectors | -2.25 | 0.25 | Inhibited | 0.45% | -1.37 |
| RHO GTPases Activate Formins | -1.97 | 0.24 | Inhibited | 0.86% | -1.24 |
| Developmental Biology | -2.20 | 0.28 | Inhibited | 0.29% | -1.21 |
| Resolution of Sister Chromatid Cohesion | -1.83 | 0.24 | Inhibited | 0.98% | -1.15 |
| Mitotic Metaphase and Anaphase | -1.97 | 0.28 | Inhibited | 0.46% | -1.08 |
| Mitotic Anaphase | -1.97 | 0.28 | Inhibited | 0.46% | -1.08 |
| Separation of Sister Chromatids | -1.46 | 0.28 | Inhibited | 0.60% | -0.80 |
| Mitochondrial biogenesis | -0.81 | 0.18 | Inhibited | 3.70% | -0.60 |
| Transcriptional activation of mitochondrial biogenesis | -0.81 | 0.18 | Inhibited | 10.00% | -0.60 |
| Classical antibody-mediated complement activation | 0.80 | 0.18 | Activated | 14.29% | 0.59 |
| Creation of C4 and C2 activators | 0.80 | 0.18 | Activated | 7.14% | 0.59 |
| Initial triggering of complement | 0.80 | 0.21 | Activated | 5.26% | 0.55 |
| Complement cascade | 0.80 | 0.21 | Activated | 2.22% | 0.55 |
| Organelle biogenesis and maintenance | -0.81 | 0.28 | Inhibited | 0.50% | -0.44 |
| Innate Immune System | 0.80 | 0.37 | Activated | 0.21% | 0.35 |

Also, key regulators of mitochondrial biogenesis, such as *Sirt3*, *Dmac2l*, *Idh2* and *Sirt5*, were among the differentially expressed genes, suggesting an inhibition of mitochondrial metabolism in long-term T3 treated mice (Fig. 4B). While most pathways were found to be inhibited, several clusters of genes associated with the function of the innate immune system suggested an activated immune response in long-term T3 treated mice after MI compared to vehicle treated WT mice after MI (Fig. 5A). Especially genes related to complement and the FcγR system were found to be upregulated in T3 treated mice, e.g. Cfd, Fcgr1, Fcgr4, C4b, Fcgr2b and C1qa. Accordingly, spleen weight (in relation to tibia length) was significantly higher in response to long-term T3 treatment in WT mice undergone MI or sham surgery (Fig 5B and 5C) compared to vehicle treated controls.

**Figure 5.**
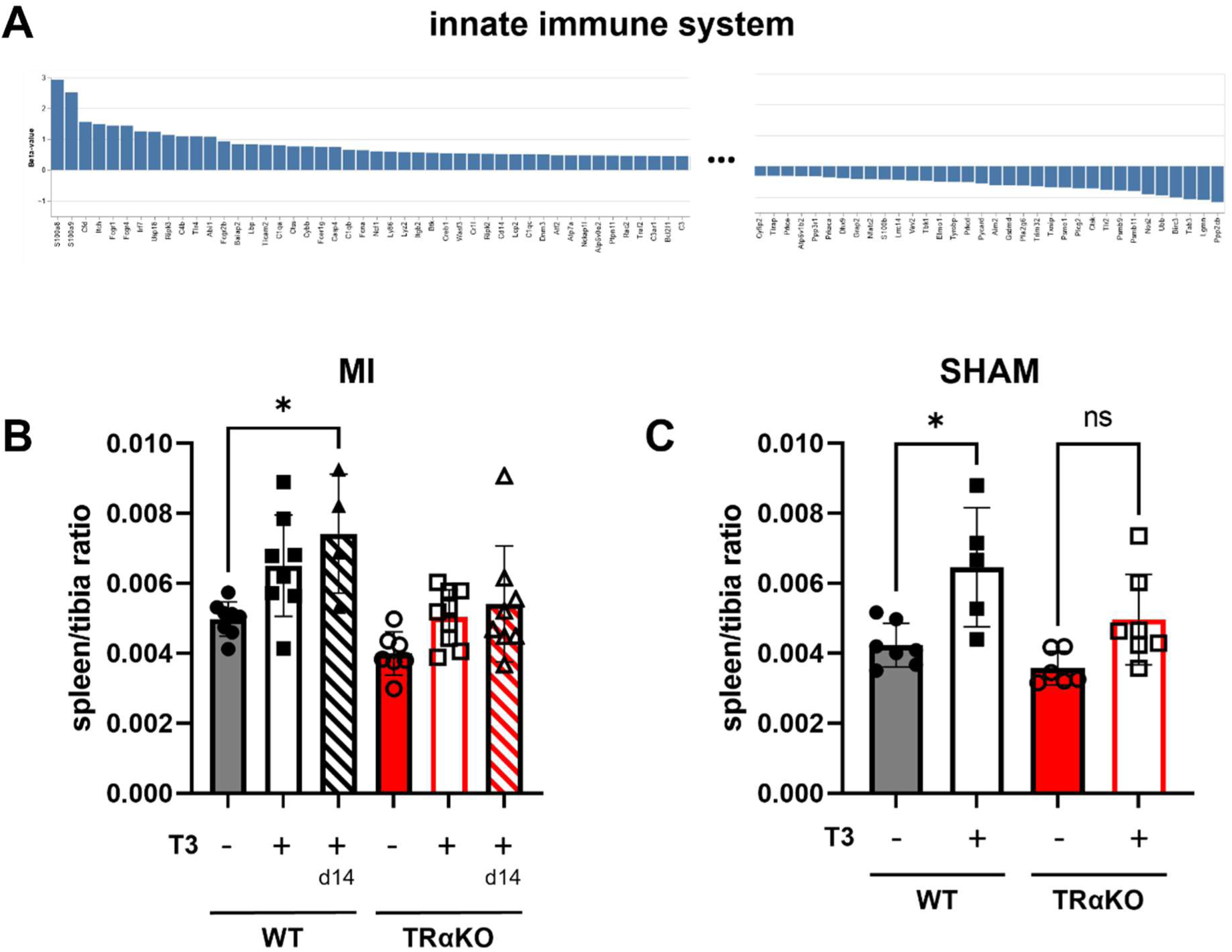
Effect of T3 administration in MI on the immune system. A) Regulated genes of the innate immune system are shown in a waterfall plot. For display purposes only the end points in this plot are shown. Spleen weight in relation to tibia length of WT and TRαKO MI (B) and SHAM mice (C). MW ± SD; *p≤0.05

## Discussion

Short-term T3 treatment for 5 days post-MI resulted in an improved cardiac function in WT mice without the induction of cardiac hypertrophy or impact on the infarct size. In contrast, long-term T3 treatment for 56 days post-MI induced cardiac hypertrophy, but failed to improve cardiac function or to reduce infarct size in WT. These data show that post-MI treatment with T3 can be beneficial, which is in line with previous reports. The comparison of short and long-term T3 treatment, however, adds important information about the correct timing and duration of T3 treatment, since prolonged T3 treatment post-MI did not reveal beneficial effects on cardiac hypertrophy or cardiac function. Altogether, beneficial and adverse effects of T3 treatment post-MI need to be considered, whereby an immediate initiation and short duration of T3 treatment appears to be crucial for improved cardiac function and protection for increased cardiac hypertrophy.

The EF of WT in the short-term T3 treatment after MI (WT MI + T3) was significantly higher compared to the vehicle treated control group (WT MI). TRαKO mice showed no difference in EF in response to short-term T3 treatment after MI, which indicates that the benefit of T3 in improving systolic function depended on TRα. The fact that T3 treatment did not increase heart weight normalized to tibia length significantly in any of the short-term treated groups, neither WT nor TRαKO relative to their respective controls, suggests that this approach does not impact on pathological myocardial remodeling and is thus rather safe.

Furthermore, the present data demonstrate that TH/TRα signaling has two distinct roles in MI, which depend on the timing: TRα absence at the time of infarction was rather protective in vehicle treated TRαKO mice compared to vehicle treated WT mice, namely the EF in vehicle treated TRαKO was less affected/reduced then in WT mice after MI. This result aligns with results from our previous ex vivo study in isolated and perfused hearts from WT and TRαKO mice (Pape et al., 2022), where the absence of TRα resulted in a smaller infarct size as assessed by TTC staining. Thus, at the time of the infarction, when oxygen supply was suddenly reduced, reduced TH signaling (TRαKO or hypothyroidism) was beneficial, possibly due to a reduction in oxygen consumption. After MI, in contrast, T3 treatment for a short time had a prolonged effect and preserved cardiac function. Thus, TH/TRα signalling appears to be adverse during infarction, but beneficial in the recovery afterwards.

Long-term T3 treatment for 56 days, however, did not result in beneficial effects. Unlike short-term T3 treatment, long-term T3 treatment led to cardiac hypertrophy and failed to improve cardiac function. In WT mice, neither immediate nor delayed long-term T3 treatment led to significant improvement in EF compared to untreated MI; thus, the concomitant cardiac hypertrophy was not an effective compensatory response.

Gene expression analyses revealed an upregulation of TRβ expression in the TRαKO hearts upon T3 treatment, which suggests that TRβ may contribute to the hypertrophic response in the absence of TRα. While this seems a likely explanation, we had previously observed that T3 induced ventricular hypertrophy in WT and TRβKO mice, but not in TRαKO mice (Geist et al., 2024). Thus, TRβ cannot not substitute for TRα. We hypothesize that an indirect effect of T3 on cardiac workload in response to MI, e.g. central stimulation of the sympathetic nervous system, that causes of the hypertrophic response to T3 long-term treatment observed in TRαKO mice post-MI within the current study.

The RNA-seq data suggested that long-term T3 treatment shifted the myocardium towards a state with enhanced contractile-cytoskeletal remodeling (possibly maladaptive), increased metabolic and mitochondrial stress or adaptation, and an activated inflammatory milieu. This triad could underlie the observed hypertrophy without functional improvement and may explain long-term T3 treatment failed to improve ejection fraction compared to control. For example, the complement and FcγR system as well as neutrophils are a known drivers of pathology during MI (Asare et al., 2025; Fang et al., 2023; Kim et al., 2024). Moreover, higher levels of Ly6G might correlate with an enhanced accumulation of Ly6G+ neutrophils in damaged tissue.

The present data have clinical implications. Improved post-MI EF after T3 treatment depended on TRα. This suggests the application of TRα-specific agonists to generate the beneficial effects while avoiding potentially adverse effects via TRβ. The same principle of improving a pathological condition with a TR agonist has recently been approved in the US and Europe for hepatic steatosis with fibrosis and TRβ with resmetirom as a hepatocyte-specific TRβ agonist (Harrison et al., 2024). In the past decade, patients with resistance to thyroid hormone due to mutations in the TRα gene (RTHα) have been reported (Bochukova et al., 2012; Moran & Chatterjee, 2015). It can be reasoned that in these patients post-MI T3 treatment may fail to induce beneficial effects. A TRα-specific agonist could be applied with higher doses and overcome the resistance without inducing general thyrotoxicosis.

In summary, short-term immediate post-MI T3 treatment improved cardiac function in WT mice without inducing cardiac hypertrophy or infarct size change. This effect depended on the presence of TRα. Long-term T3 treatment for 56 days induced hypertrophy but failed to improve function or reduce infarct size in WT. To use the beneficial effects of T3 in MI, these data suggest that treatment needs to start early and should be continued only for a limited time.

## Acknowledgments

We are grateful for the continued dedicated support from Prof. Dr. G. Hilken, Dr. A. Wißmann, Dr. P. Dammann, Dr. M. Dubicanac and the staff of the animal facility at the University Hospital Essen. We thank Konstanze Schättel, Nadine Yurdagül-Hemmrich, Ann-Kathrin Weiss and Kristina Piwellek for their dedicated technical assistance. We thank Bettina Budeus for expert support with RNA-Sequencing.

